# Inoculation with *Pisolithus tinctorius* may ameliorate acid rain impacts on soil microbial communities associated with *Pinus massoniana* seedlings

**DOI:** 10.1101/416834

**Authors:** Mia R. Maltz, Zhan Chen, Jixin Cao, Keshav Arogyaswamy, Hannah Shulman, Emma L. Aronson

**Author notes:** Co-first Authors, denotes equal author contributions. Corresponding Author: Zhan Chen Please direct email correspondence to (MM), incare of Zhan Chen; (MM); Website: http://sites.uci.edu/maltz/ (MM) Submitted for publication in Fungal Ecology in 2018.

## Abstract

Human activities accelerate acidification, particularly as acid rain, which may have lasting impacts on soil abiotic and biotic parameters. However, the effects of acidification on aboveground vegetation, belowground communities, and carbon cycling remains unresolved. We examined the effects of long-term acidic treatments and *Pisolithus* inoculation on plants, soils, and microbial communities in pine plantations and found that exposure to severely-acidic treatments diminished plant performance, altered microbial communities, and decreased organic matter, nitrate, and available phosphorus. Although we did not detect any benefits of *Pisolithus* inoculation for *Pinus* seedlings impacted by severely-acidic treatments, when these severe treatments were inoculated with *Pisolithus*, both soil properties and microbial community composition shifted. We posit that inoculation with *Pisolithus* may alleviate stressful environmental conditions, and change the structure of mycorrhizal fungal communities. Although acidification may alter biogeochemical cycles and constrain aboveground and belowground communities, *Pisolithus* inoculation may provide benefits to some components of forested ecosystems.

## Introduction

Microbes play a critical role in biogeochemical cycling as regulators of decomposition, plant-microbe interactions, and nutrient transformations (van der Heijden et al. 2008). However, microbial processes are threatened by human activities and industrial inputs to ecosystems, such as sulfur (S) and nitrogen (N) deposition, which result in landscape degradation and biodiversity loss (Bobbink et al. 2010, Allen et al. 2016). Industrial inputs can acidify soils, further threatening microbial processes and ecosystem functions. Accelerated acidification in forest soils via acid deposition from anthropogenic sources may have lasting impacts on soil physicochemical properties and biotic communities. For example, deposition from industrial emissions on forests in Europe (Jandl et al. 2012) and North America (Reininger et al. 2011) have been linked to long-lasting disruptions in biotic communities and biogeochemical cycling (Likens et al. 1996). However, the long-term effects and the degree to which industrial inputs leading to soil acidification impacts belowground carbon cycling via shifts in plant and soil microbial community composition in terrestrial ecosystems is not yet resolved.

While variable, acid rain generally reduces plant productivity by damaging foliar tissue (Neufeld et al. 1985), reducing biomass (Raschke 1975), diminishing resource availability to plants (Bååth & Arnebrandt 1994), and disrupting essential physiological processes including stomatal functioning (Zieger et al. 1983) and chlorophyll production (Odiyi and Bamidele 2013). Acid deposition dissolves nutrients and divalent cations, such as magnesium (Mg) and calcium (Ca) (Berger et al. 2016, Leys et al. 2016, Türtscher et al. 2017), while facilitating aluminum (Al) release which could reduce plant root growth and inhibit plant nutrient uptake (Burnham et al. 2017). By stripping soils of nutrients, hindering soils’ capacity to buffer plants from toxic substances, and physically harming aboveground vegetation, acid deposition can reduce plant productivity through malnourishment, dehydration (Macaulay et al. 2015), or toxicity.

By constraining plant productivity, acidification could indirectly alter the composition of fungal communities and hamper soil microbial processes, with implications for organic carbon storage in forested ecosystems (Kemmitt et al. 2006, Janssens et al. 2010, Oulehle et al. 2011). Even though the consequences of these inputs on soil organic matter (SOM) content have been inconsistent (Averill & Waring 2017, Liu et al. 2011, Mack et al. 2004), reductions in microbial CO_2_ efflux and soil organic carbon turnover associated with soil acidification could lead to SOM accumulation (Tonneijic et al. 2010) by inhibiting decomposition of woody debris (Ferreira et al. 2017), and by influencing microbial physiology (Chen et al. 2016), growth (Rousk et al. 2010), and activity (Anderson & Domsch 1993).

Through damaging plant communities, acidification in soil could also interfere with the functioning of plant symbionts, such as mycorrhizal fungi (Carrino-Kyker et al. 2016). Ectomycorrhizal fungi (EM) play essential roles in forested ecosystems (McNear Jr. 2013), such as soil exploration and mineral weathering, and by acting as a link between aboveground and belowground components of biogeochemical cycles (Treseder & Allen 2000). A subset of EM fungi produces extracellular enzymes (Feng 1993, Bodeker et al. 2009), which amplify the bioavailability of rhizosphere nutrients (Courty et al. 2005, 2010a, Talbot et al. 2015). Because EM fungi obtain a majority of their C from live plants, mycorrhizal biomass and resource allocation patterns are largely reliant on the vitality of plant foliage. Previous studies have shown that acidifying pollutants extensively damage forest tree roots and EM fungi (Sobotka 1964, Stroo and Alexander 1985), via increased metal toxicity or foliar deterioration. These effects likely limit photosynthetic capacity, carbohydrate flux to roots, and mycorrhizal function (Dighton & Jansen 1991). As soil pH plays an important role in regulating EM turnover (Glassman et al. 2017, Molina & Trappe 1982), acid-induced changes in soil chemistry can also directly alter EM community structure (Markkola & Ohtonen 1998), diversity (Kluber et al. 2012), and hyphal length (Dighton & Skeffington 1987). Although EM fungi aid their host plants in efficiently acquiring nutrients (Nara et al. 2003, Qu et al. 2010, Chorianopoulou et al. 2015), acidic inputs that reduce plant diversity (Bobbink et al. 2010) or alter plant performance may also indirectly reduce ectomycorrhizal growth and EM root colonization (Arnolds 1988, Wallenda & Kottke 1998).

Along with EM fungal communities and functions, other microbial groups may also be sensitive to the effects of acidification. The diversity and relative abundances of some bacterial taxa, such as *Acidobacteria, Actinobacteria,* and *Bacteriodetes,* change predictably across soil acidity gradients (Jones et al. 2009). Although a number of bacterial taxa may be excluded from acidic environments, such as acid-impacted lakes, several bacterial taxa, including Alphaproteobacteria may thrive at low pH (Percent et al. 2008). Previous studies have shown that pH is an accurate predictor of soil bacterial community structure across large spatial scales (Lauber et al. 2009, Landesman et al. 2014). In their biogeographic survey across the Western Hemisphere, Fierer and Jackson (2006), found that the most acidic soils in the Peruvian Amazon had lowest bacterial diversity. In a biogeographic assessment across Great Britain, Griffiths et al. (2011) similarly showed a positive relationship between alpha diversity and soil pH; conversely, in the same study, beta diversity was greatest in acidic soils.

Saprotrophic fungal communities may differ as soil and plant-litter chemistries change (Prewitt et al. 2014). Surveys of macrofungal fruitbodies showed that ratios of saprotrophic to mycorrhizal fungi tend to increase when forests decline in productivity (Fellner & Pešková 1995). Factors that reduce forest productivity could either reduce the relative abundance of mycorrhizal fungi or increase the percentage of decomposer fungi in the fungal community, with implications for forest nutrient cycling.

As ecosystems recover from long-term effects of acidification, both aboveground and belowground components of ecosystems may benefit from active mycorrhizal restoration via EM inoculation to augment revegetation efforts (Leake 2001). Inoculation with EM fungi may improve tree performance and promote ecosystem recovery from long-term acidification. Some EM, such as *Pisolithus,* grow well in acidic environments, like acidic strip mine sites (Hendrix et al. 1985, Liang et al.1999, Baroglio et al. 2000). Mycorrhizal fungi may not only assist host trees in acquiring macronutrients (Virant-Klan & Gogala 1995, Chuyong et al. 2000), but may also buffer plants from the effects of acid rain. However, little is known about the levels of soil acidity which effectively diminish the capacity of EM fungi to associate with or provide benefits to their plant hosts. Although belowground communities may predictably respond to acidic inputs, direct restoration of soil microbes and plant-associated fungi may foster ecosystem succession towards alternative stable states within recently restored systems (Lewontin 1969, Rustad et al. 1996, Bardhan et al. 2012). An explicit consideration of both fungal and bacterial communities may provide valuable metrics for directing conservation efforts and assessing soil ecosystem recovery from acidification in forested ecosystems (Baldrian et al. 2012).

Acid rain inputs and soil acidification are major concerns in Asia (Zhao et al. 2009, Yang et al. 2012). China is a prime example of acidification because most ecosystems in southern China receive large quantities of acidic inputs (Zhang et al. 2012). In southern China, acid rain has detrimentally affected plant and soil traits (e.g., Fan & Wang 2000, Wang et al. 2009, Zhang et al. 2007, Lv et al. 2014). Yet, little is known about the long-term impacts of acid rain on soil microbial communities and biogeochemical cycling in forested ecosystems, or whether microbial inoculation ameliorates these effects on Chinese forests.

*Pinus massoniana* is a widespread native pine species in central and southern China. To compensate for the loss of natural forests in southern China, *P. massoniana* is commonly grown in pine plantations. This tree forms EM associations with a number of EM fungal taxa. However, *P. massoniana* is sensitive to acid rain (Yu et al. 2017) and may benefit from strategies lessening the impact of acidification on pine forests. Our previous experiments in southern China showed that inoculation with the EM taxon *Pisolithus tinctorius* may ameliorate the effects of acidification and improve the growth of *Pinus massoniana* seedlings under different acid rain treatments (Chen et al. 2013, Chen & Shang 2014).

To examine the long-term effects of acid rain treatments on plant and soil properties and microbial communities and to evaluate the extent to which these effects are ameliorated by ectomycorrhizal inoculation in forested ecosystems in southern China, we experimentally addressed the following questions: (1) Does acid rain have inhibitory effects on plant, soil, and microbial communities? And (2), does active mycorrhizal restoration increase plant growth, improve soil properties, and promote microbial diversity following protracted acid rain exposure? To approach these questions, we first investigated the effects of acidic treatments on plant performance in a forested ecosystem in southern China by evaluating soil properties, nutrient concentrations, and soil organic matter in both acid-treated and inoculated treatment plots. Next, we examined fungal and bacterial communities to determine whether *Pisolithus* inoculation ameliorated the effects of acid treatments on soil microbial communities and plant hosts. We hypothesized that in acidic treated *P. massoniana* plots (1) plant performance would decline, (2) both soil nutrients and soil organic matter would decrease, and (3) soil microbial communities would be altered. If acidification reduces the abundance or functioning of EM fungi, then *P. massoniana* seedlings may struggle to persist in pine plantations without fungal mutualists. Therefore, we hypothesized that (4) inoculation with *Pisolithus* would facilitate better tree growth and resilience than acid-treated trees without *Pisolithus*, along with associated changes to soil properties, biogeochemical cycling, and microbial communities in acid-treated *Pisolithus* inoculated plots.

## Materials and Methods

### Study system

We investigated the effects of acidic treatments on plants, soils, and microbial communities in experimental plots planted with *Pinus massoniana* in Changsha County, Hunan Province (N28°26′9.13″ E113°11′39.26″). The mean annual temperature is 17.2°C and mean annual precipitation at our study site is 1422.4 mm. This location is frequently exposed to acidic precipitation, with a mean annual pH of precipitation of 5.02 (Environmental Quality of Hunan Province in 2016 by Environmental Protection Department of Hunan, hnhbt.hunan.gov.cn). Prior to treatment, the pH of field soils at our experimental site were 5.73 0.13 (Table 1, Supplemental Table SI1).

**Table 1.** Acidic solution and soil pH across time and among treatment combinations.

| pH solution<br>+/- *P.t.* (*Pisolithus*<br>inoculation) | pH soil<br>2015<br>Pre-treatment | pH soil<br>2015 | pH soil<br>2016 |
| --- | --- | --- | --- |
| Severe<br>3.5 + *P.t.* | 5.03 ± 0.08 | 5.46 ± 0.14 | 5.57 ± 0.09 |
| Severe<br>3.5 – | 5.92 ± 0.08 | 5.90 ± 0.13 | 6.80 ± 0.09 |
| Moderate<br>4.5 + *P.t.* | 5.39 ± 0.04 | 5.71 ± 0.09 | 6.64 ± 0.003 |
| Moderate<br>4.5 – | 6.17 ± 0.12 | 6.03 ± 0.06 | 6.65 ± 0.04 |
| Ambient<br>5.6 + *P.t.* | 6.47 ± 0.38 | 6.0 ± 0.14 | 6.69 ± 0.04 |
| Ambient<br>5.6 – | 5.41 ± 0.17 | 5.43 ± 0.39 | 5.84 ± 0.13 |

Our experimental plots were located within a cleared field which was previously cultivated, but allowed to lay fallow for four years prior to our study. Prior to initiating our experimental treatments, in March 2015 we analyzed baseline soil nutrients within our field plots and found that soils contained total N of 0.13 g kg^-1^, Olsen – phosphorus (P) of 28.59 mg kg^-1^, and an available potassium (K) of 31.5 mg kg^-1^. Using a total organic carbon (TOC) analyzer, we found that soil organic matter was 20.28 g kg^-1^. We constructed eighteen 6 m^2^ experimental plots; each plot was covered with a canopy, which extended 5 m above the plots. This canopy enclosed our experimental plots, thus preventing natural precipitation from directly reaching soil surfaces. Outside of each treatment plot, we left an untreated 0.5 m – 0.6 m gap surrounding each edge of each treatment plot.

#### Inoculation treatment

In March 2013, *Pinus massoniana* seedlings were inoculated with either *Pisolithus tinctorius* EM fungal inoculum (*Pisolithus tinctorus* [Pers.] Coker et Couch; Sclerodermatacea) or sterilized *P. tinctorius* material. *Pisolithus tinctorus* was isolated from Jiangxi Province and cultured in vermiculite and peat. We autoclaved *P. tinctorius* material for one hour at 0.11MPa pressure and 121°C to eliminate any viable *Pisolithus tinctorus* spores or hyphal fragments in the sterilized material.

Both inoculated and uninoculated *P. massoniana* seedlings were grown in natural light under a canopy of transparent film, and watered, as needed, for two years prior to outplanting. In March 2015, 30 inoculated *P. massoniana* or 30 *P. massoniana* uninoculated seedlings (with autoclaved inoculum added) were planted in each experimental plot. Seedlings were maintained in our experimental plots for 20 months.

#### Acidic treatment

Beneath the constructed canopy, we treated soils in our experimental plots with solutions consisting of either a moderately-acidic (pH 4.5) solution or a severely-acidic (pH 3.5) solution, or an ambient treatment (pH 5.6); ambient treatments received solutions under the same constructed canopy. Each experimental treatment plot was replicated three times per treatment combination, totaling 18 experimental plots. To achieve pH treatment solutions, deionized water was treated with an acidic solution composed of sulfuric acid (H_2_SO_4_ ^2-^) and nitric acid (HNO ^-^), to lower the pH of the solution to either pH 4.5 (moderately-acidic) or pH 3.5 (severely-acidic), or for ambient solutions to reach a pH of 5.6. No additional nutrients were added to our solutions. Each *P. massoniana* seedling, including those in ambient treatments, received 6.7 L of solution per week, which corresponded to 1.1 L m^-2^ per week, for 20 months. Ambient treatments received the same solutions as the severely and moderately-acidic treatments, except the pH of ambient treatments was 5.6, which was more basic than the mean annual pH of precipitation in the region. The mole ratio of SO_4_^2-^ to NO_3_^-^ in the acidic solutions was 4:1. Other ion concentrations in the solutions were NH_4_^+^ - 2.67 mg L^-1^, Ca^2+^ - 3.37 mg L^-1^, Mg^2+^ - 0.33 mg L^-1^, Cl^-^ −14 mg L^-1^, K^+^ - 0.79 mg L^-1^, Na^+-^ 0.36 mg L^-1^, F^-^ - 0.39 mg L^-1^. In addition to spraying treatment solutions on seedlings in experimental plots, plots were consistently watered with local groundwater, pH 6.2, up to several times weekly, as needed, throughout the duration of the experiment to make up for the deficit between the amount of treatment water under the canopy and the ambient rainfall. Amounts applied to plots varied by ambient seasonal precipitation patterns to eliminate drought stress, and did not vary by treatment or plot.

### Sample collection

Plants and soil samples were collected in both November 2015 and November 2016, after 8 and 20 months, respectively. Three randomly selected seedlings were destructively sampled per experimental plot, and plant height, diameter, and fresh weight biomass was recorded from each sample. Plant samples were then oven dried at 60 °C for 48 h, and weighed on an analytical balance to determine the dry weight biomass. We recorded total plant biomass, aboveground biomass, belowground biomass, and needle weights from each individual, as well as soil properties (SI Table 2-SI Table 3).

**Table 2.** *Pinus massoniana* properties in response to inoculation and acidic treatments.

| pH solution<br>+/- *P.t.*<br>(*Pisolithus*<br>inoculation) | Plant<br>diameter<br>(cm) | Plant height<br>(cm) | Plant<br>root:shoot | Dried stem<br>biomass (g) | Dried needle<br>biomass (g) |
| --- | --- | --- | --- | --- | --- |
| Severe<br>3.5 + *P.t.* | 15.37 ± 0.56<br>(b,c) | 116.67 ± 1.67<br>(e) | 14.31 ± 2.20<br>(f) | 66.98 ± 5.42<br>(b,c) | 73.29 ± 5.63<br>(b) |
| Severe<br>3.5 - | 13.75 ± 0.58<br>(c) | 117.33 ± 3.93<br>(e) | 12.62 ± 0.32<br>(f,g) | 60.35 ± 7.94<br>(c) | 71.88 ± 13.86<br>(b) |
| Moderate<br>4.5 + *P.t.* | 18.40 ± 0.56<br>(a) | 149.33 ± 3.84<br>(d) | 13.06 ± 1.24<br>(f,g) | 135.07 ± 8.74<br>(a) | 142.55 ± 13.08<br>(a) |
| Moderate<br>4.5 - | 15.99 ± 1.43<br>(b,c) | 142.00 ± 4.16<br>(d) | 12.52 ± 0.21<br>(f,g) | 80.98 ± 14.56<br>(b,c) | 154.25 ± 5.20<br>(a) |
| Ambient<br>+ *P.t.* | 15.97 ± 0.36<br>(b,c) | 156.33 ±<br>10.93 (d) | 11.23 ± 0.61<br>(f,g) | 123.77 ± 9.59<br>(a) | 136.58 ± 4.16<br>(a) |
| Ambient - | 17.60 ± 0.37<br>(a,b) | 147.67 ± 6.74<br>(d) | 10.58 ± 0.35<br>(g) | 90.21 ± 6.67<br>(b) | 129.77 ± 6.87<br>(a) |

Three soil cores (20 cm x 3.4 cm) were randomly collected from within the drip line (under the canopy) of treated plants within each plot. Each sample was homogenized and subsequently used to analyze soil properties. Soil subsamples (100 g per subsample) were immediately frozen on dry ice for transport back to the laboratory, and stored at −80°C for molecular analyses of microbial communities. We weighed 0.25 g aliquots of each frozen soil sample in triplicate to assay for microbial community analyses. Next, we extracted DNA from these soil aliquots and targeted hypervariable portions of both the V3–V4 region of the bacterial 16S rRNA gene and the fungal internal transcribed spacer (fungal ITS1) for Polymerase chain reactions (PCR) amplification. We purified and quantified our PCR products, prior to pooling in equimolar concentrations for Illumina Miseq sequencing (Illumina Incorporated, California; SI Molecular Methods).

### Statistical analysis

To test our hypothesis that plant performance would decline in acidic treated *P. massoniana* plots, we performed a series of analyses of variance (ANOVAs) with acidic solution treatment as the independent variable and plant height or plant biomass as the dependent variables. To address our second hypothesis, we examined the response of soil organic matter and soil nutrients to acidic treatments by performing response ratios. In each of these cases, the independent variable was acid treatment level, and the dependent variable was soil organic matter or soil nutrients. The ratio of the values for each treatment combination was log_2_-transformed to provide a response factor. To disentangle the direct effects of acid treatments on plants with the indirect effects of acid treatment on mycorrhizal fungi, we measured the effect of *Pisolithus* inoculation on *P. massoniana*. Where indicated, Tukey post-hoc tests for pairwise comparisons were applied.

We constructed a linear model to address our third and fourth hypotheses that acid treatments would alter the microbial community and *Pisolithus* inoculation promotes plant, soil, and microbial recovery. For examining microbial and fungal community shifts and to determine the relationship between fungal or microbial community composition and acidic treatments, we used a PERMANOVA analysis of microbial community data using Bray–Curtis dissimilarity and the adonis function in the vegan package of R (Anderson et al. 2008; Oksanen et al. 2012). In this model, we investigated the relationship between fungal and microbial community composition and inoculation in both 2015 and 2016. In order to account for plot to plot variation, we used ‘strata’ in adonis2 to restrict permutations solely within our spatial variable (i.e., row proximity to the Jingouba River). We examined how fungal and bacterial community composition shifted in relation to soil nutrients and physicochemical properties, including the ratio of divalent cations to total free Al, by including these environmental variables in our PERMANOVA analyses.

Principal coordinate analyses (PCoA) plots with Bray-Curtis dissimilarity metrics were used to visualize fungal and bacterial community composition, and to determine whether inoculation was related to microbial community composition or response to acidic treatments. We bioinformatically examined differences in fungal functional groups, trophic levels, and guilds through the use of the FUNguild database (Nguyen et al. 2016) and conducted a series of general linear models to examine shifts in fungal functional groups among our treatment groups. Significant shifts in microbial community composition, increases in soil organic matter, or increased similarity to ambient treatments along with inoculation would suggest that microbial community responses may be related to recovery from the long-term effects of acidification.

## Results

### Plant performance

After two years of treatments, shoot heights and biomass of severely-acid treated plants were lower than untreated plants (p<0.05) (Table 2). Specifically, we observed that both dried stem biomass and dried needle biomass were significantly lower in plants from severely-acidic treated plots (p<0.05). Moderately-acidic treatments had no effect on plant growth (p>0.05). Uninoculated severely-acid treated plants were shorter, had lower stem and needle biomass, and had significantly narrower diameter than uninoculated ambient seedlings (p<0.05). Inoculation with *Pisolithus* increased plant diameter and dried stem biomass (p<0.05), but only in the moderately-acidic treatments.

### Soil nutrients

Soil organic matter content in severely-acid treated plots was significantly lower in uninoculated treatments than in all other treatments (p<0.05) (Fig 1). However, when severely-acid treated plots were inoculated with *Pisolithus*, SOM increased to levels equivalent to ambient plots. Although severely-acid treated plots had lower available P, K, NH_4_-N, and NO_3_-N than ambient plots, the inoculated plots treated with severely-acidic treatments had significantly higher P, K, NH_4_-N, and NO_3_-N than the uninoculated plots treated with the same severely-acidic solution (SI Table 1, Figs 2-4).

**Fig 1.**
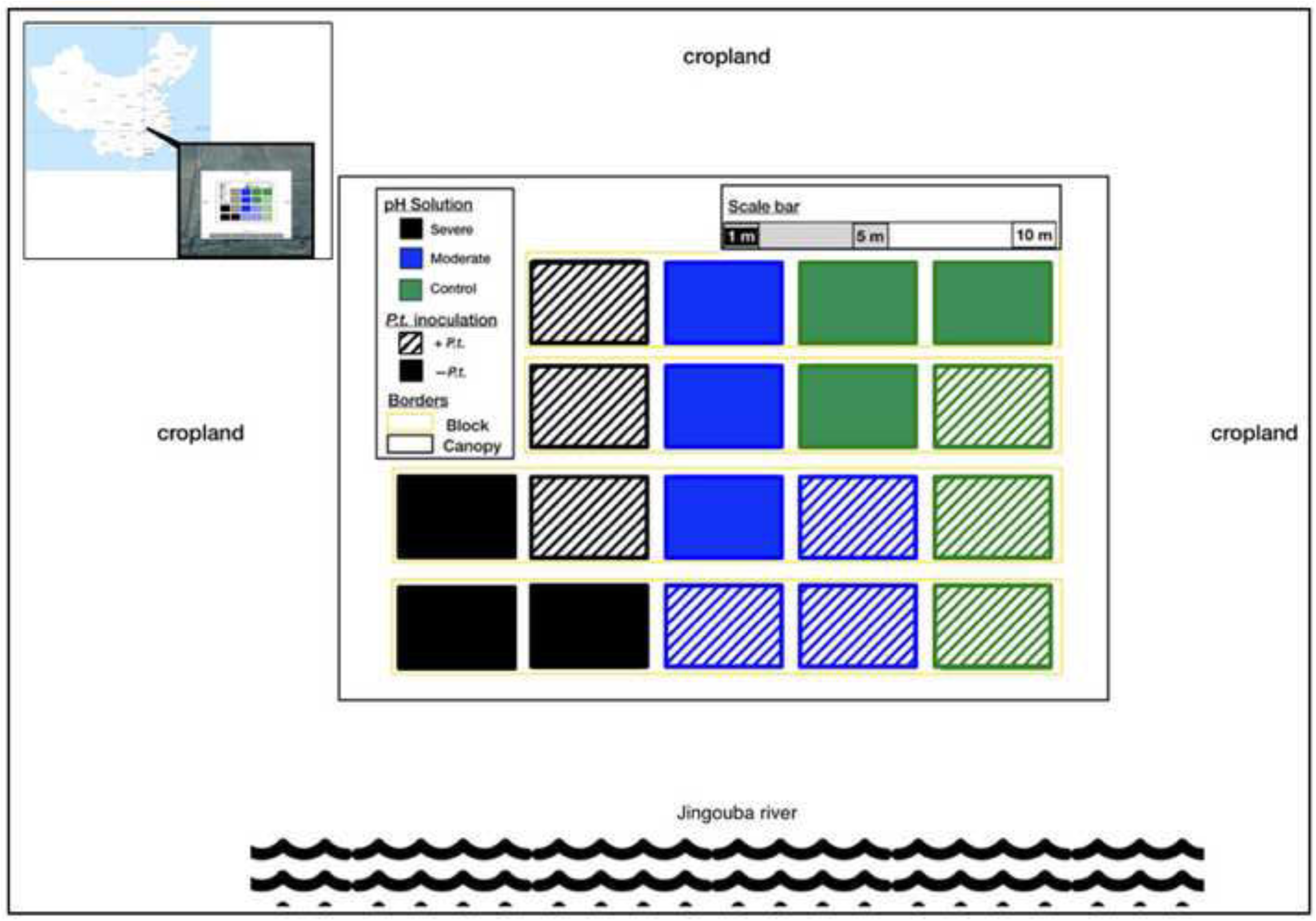
Map of the study system with scale bar. Location of our experimental plots within the Changsha County, Hunan Province (N28°26′9.13″ E113°11′39.26″). Acidic treatment solutions are colored in black for severely acidic treatment, blue for moderately acidic treatment, and green for ambient treatment. Treatment plots inoculated with *Pisolithus tinctorius* have patterned fill, while uninoculated treatments have solid fill. Black outline denotes footprint of covered canopy, yellow borders indicate row from Jingouba River, as a spatial delimiter within the study system. Inset map of China indicates where the experimental plots are located geographically.

**Fig 2.**
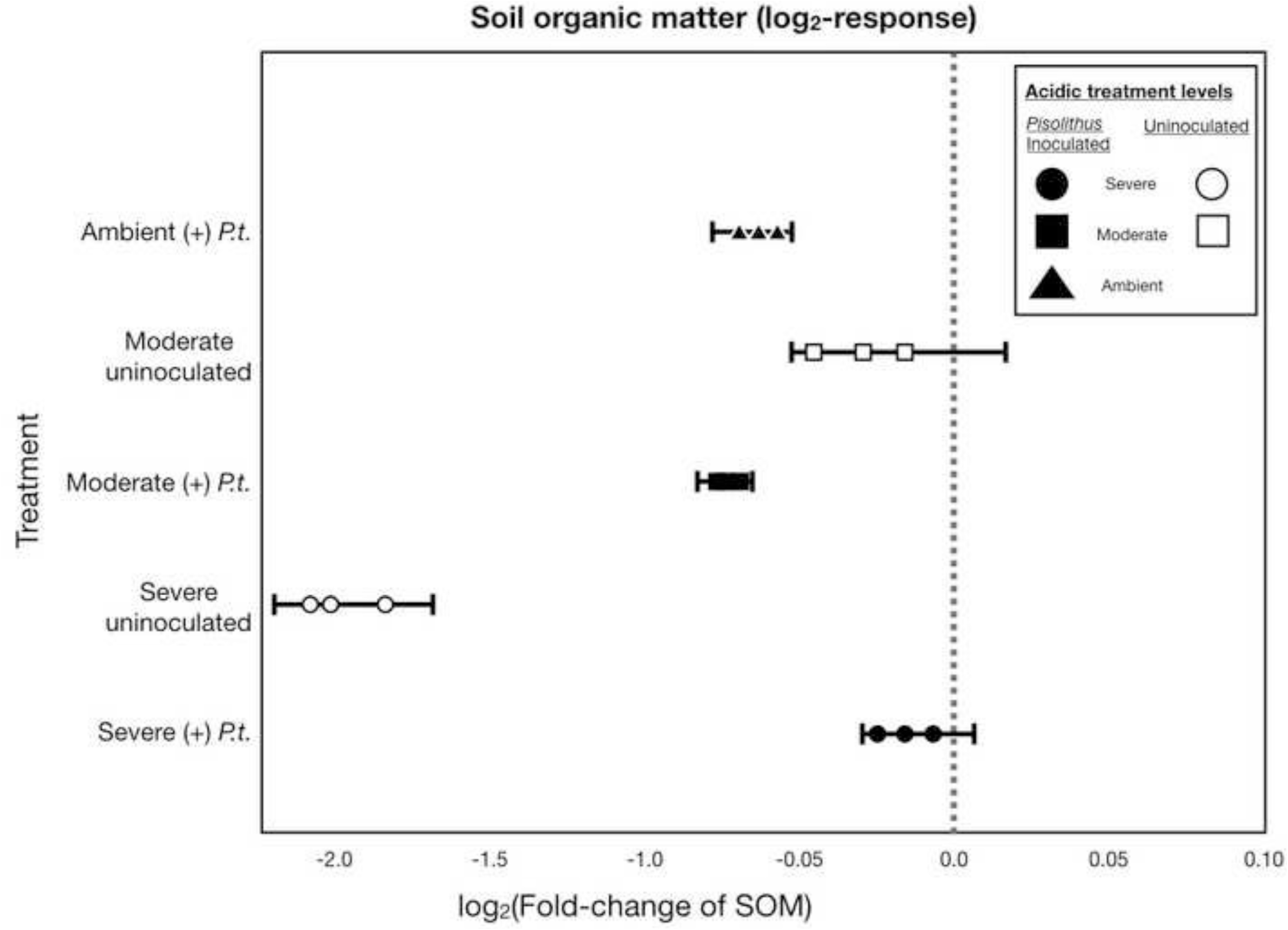
Response of soil organic matter to acidic treatments and *Pisolithus* inoculation. Response of organic matter content in g kg^-1^ of soil to acid and *Pisolithus* inoculation treatments, as compared to the organic matter content of uninoculated ambient plots. Responses log_2_ transformed, thus each unit on the horizontal x-axis represents a two-fold change. Error bars correspond to 95% confidence intervals of the mean. If points and error bars cross the dashed line at 0.0, it indicates that the particular treatment is statistically equivalent to the ambient. If points and error bars are completely below 0.0 that indicates that soil organic matter is significantly less than (left of) the untreated ambient plots; if points and error bars are completely above (to the right of) 0.0, that would indicates that soil organic matter significantly increased in that particular treatment, as compared to untreated ambient plots.

### Microbial communities

When controlling for the effect of plot location by restricting our PERMANOVA analyses by proximity to the Jingouba River (our spatial variable) to account for any spatial heterogeneity in our study site, we detected significant main effects of acidic treatment solution and inoculation for fungal communities. We found that variation in fungal communities was determined by acidic treatment solution in both 2015 (p=0.003) and 2016 (p<0.001), and inoculation in 2015 (p=0.009) and 2016 (p=0.008). Also for fungal communities, we detected significant interactions between pH of treatment solution and inoculation in 2015 (p=0 .017) and in 2016 (p<0.001). For bacterial communities, we determined that there were significant main effects of acidic treatments in both 2015 (p=0.017) and 2016 (p=0.049) and *Pisolithus* inoculation in 2015 (p=0.009) and 2016 (p=0.031), as well as significant interactions among acidic treatment and inoculation in 2015 (p<0.001) and 2016 (p<0.001).

After two years, fungal and microbial communities differed by acidic treatments (Fig 5 and 6), with distinct fungal communities from each treatment combination. Differences in fungal communities were more pronounced in 2016, which represented the longest duration of time, than they were in 2015. The PCoA Axis 2 in both 2015 and 2016 represented ∼ 17% of the variation in fungal communities. PCoA Axis 1 in 2015 represented 51.5% and Axis 1 in 2016 represented 22.6 % of the variation in fungal communities (Fig 4). No sequences from *Pisolithus* were detected in any of the treatments with autoclaved *Pisolithus* added. Fungal communities in *Pisolithus* inoculated treatments were significantly different than treatment groups with autoclaved *Pisolithus* added (p < 0.001).

**Fig 3.**
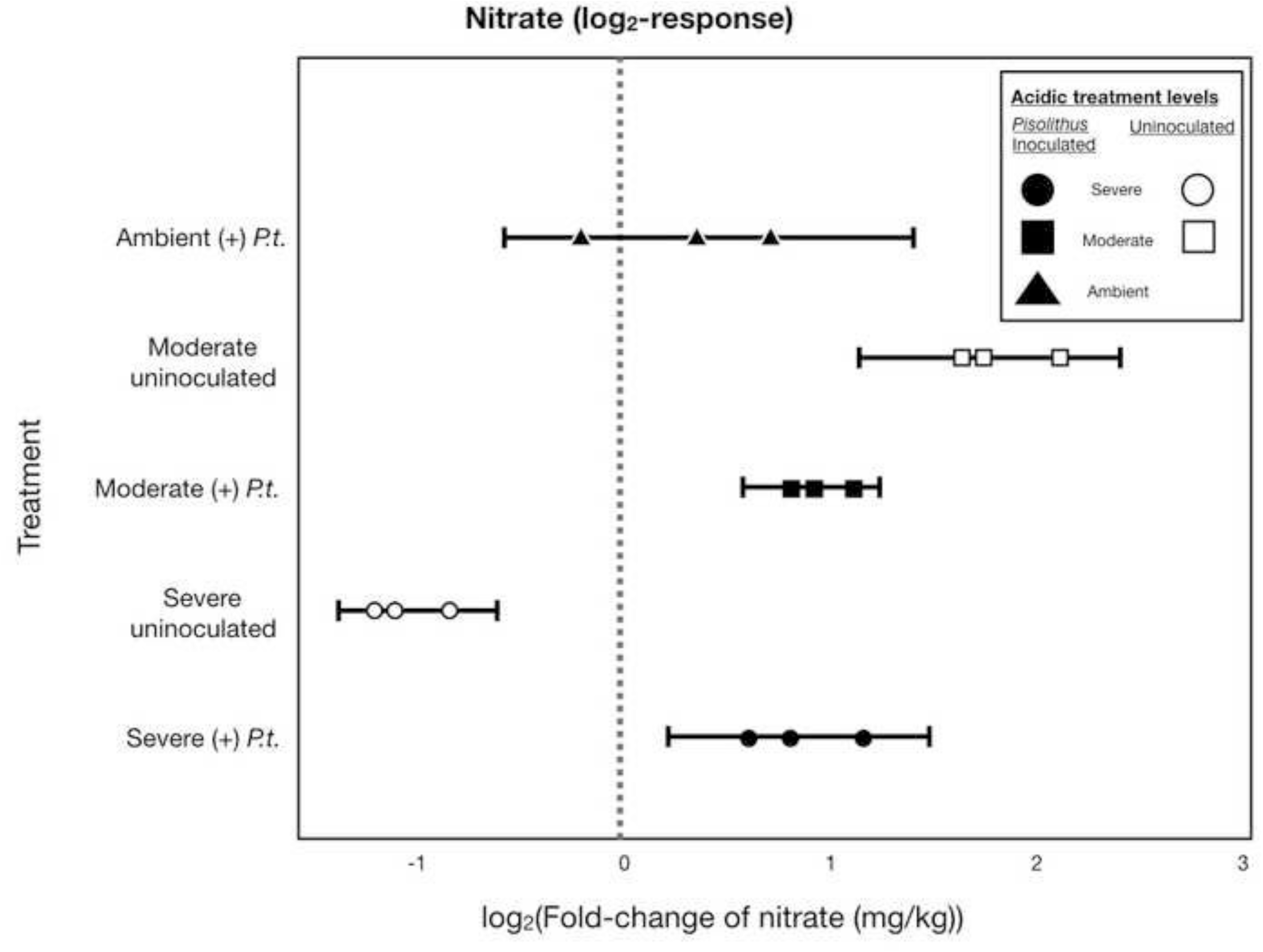
Response of nitrate to acidic treatments and *Pisolithus* inoculation. Response of nitrate in mg/kg of soil to acid and *Pisolithus* inoculation treatments, as compared to the organic matter content of uninoculated untreated ambient plots. Responses log_2_ transformed, thus each unit on the horizontal x-axis represents a two-fold change. Error bars correspond to 95% confidence intervals of the mean. If points and error bars cross the dashed line at 0.0, it indicates that the particular treatment is statistically equivalent to the ambient. If points and error bars are completely below 0.0 that indicates that soil nitrate (mg kg^-1^) is significantly less than (left of) the untreated ambient; if points and error bars are completely above (to the right of) 0.0, that would indicates that soil nitrate significantly increased in that treatment, as compared to untreated ambient plots.

**Fig 4.**
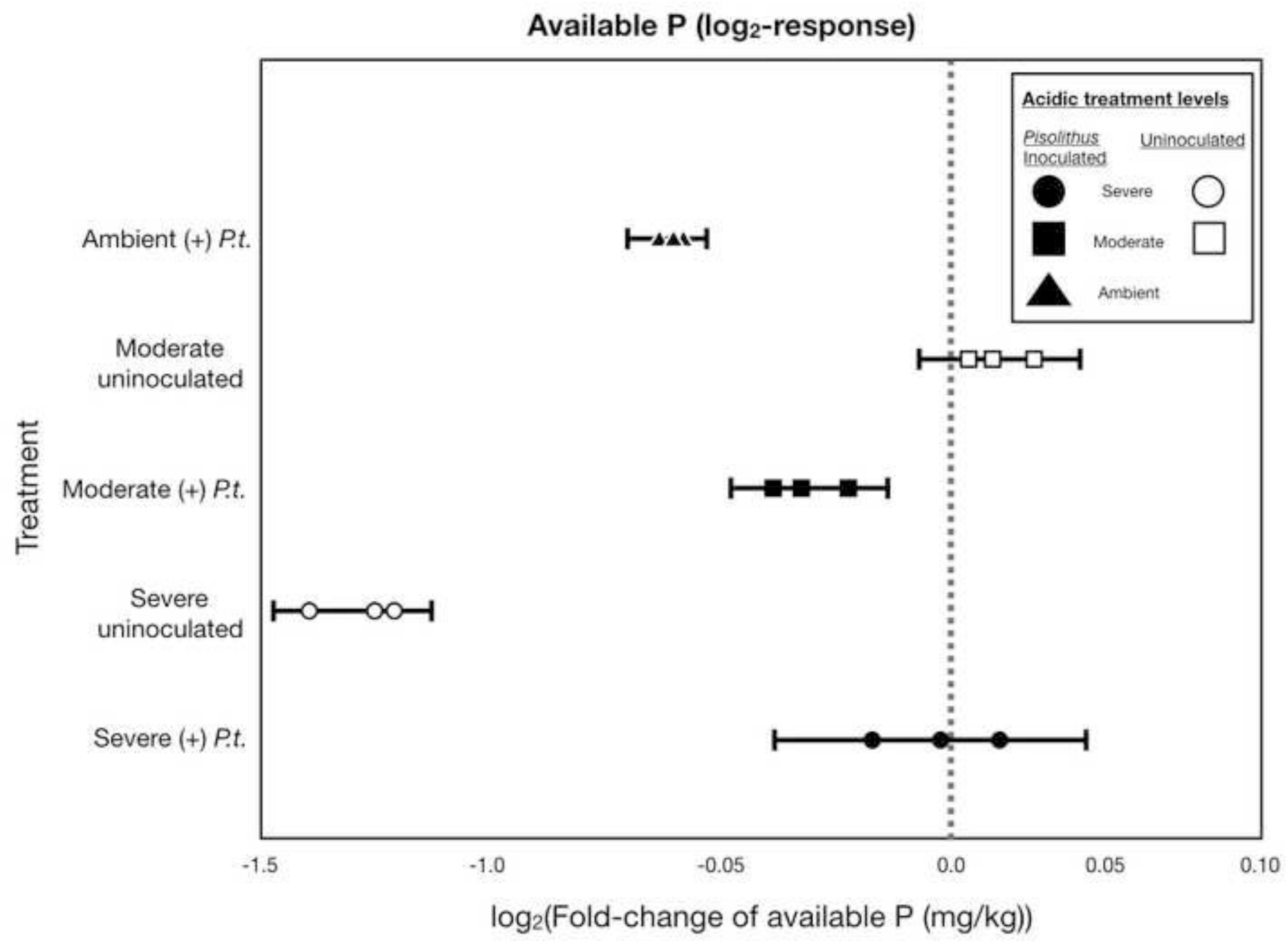
Response of available phosphorus to acidic treatments and *Pisolithus* inoculation. Response of available phosphorus (mg kg^-1^) to acid and mycorrhizal inoculation treatments, as compared to the available phosphorus in uninoculated ambient plots. Responses log_2_ transformed, thus each unit on the horizontal x-axis represents a two-fold change. Error bars correspond to 95% confidence intervals of the mean. If points and error bars cross the dashed line at 0.0, it indicates that the particular treatment is statistically equivalent to the ambient. If points and error bars are completely below 0.0 that indicates that available phosphorus is significantly less than (left of) the untreated ambient; if points and error bars are completely above (to the right of) 0.0, that would indicates that available phosphorus significantly increased in that particular treatment, as compared to untreated ambient plots.

**Fig 5.**
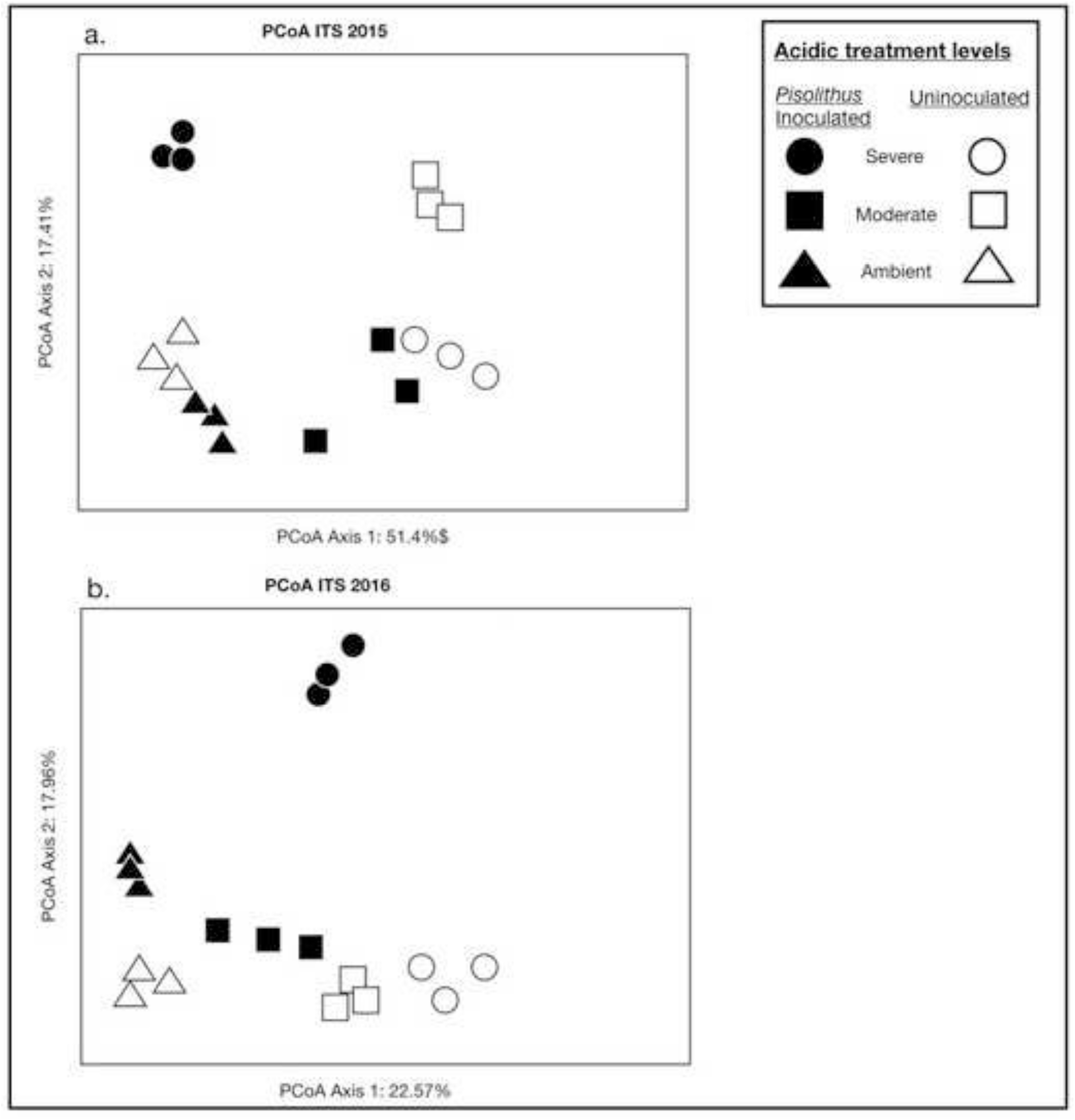
Principal coordinate analyses of fungal communities. Principal coordinate analysis (PCoA) plot of fungal ITS1 sequencing results from 2015 (a) and 2016 (b), showing fungal community responses to acid and *Pisolithus* inoculation treatments. Solid shapes indicated inoculation with *Pisolithus*; open shapes indicate uninoculated treatments.

**Fig 6.**
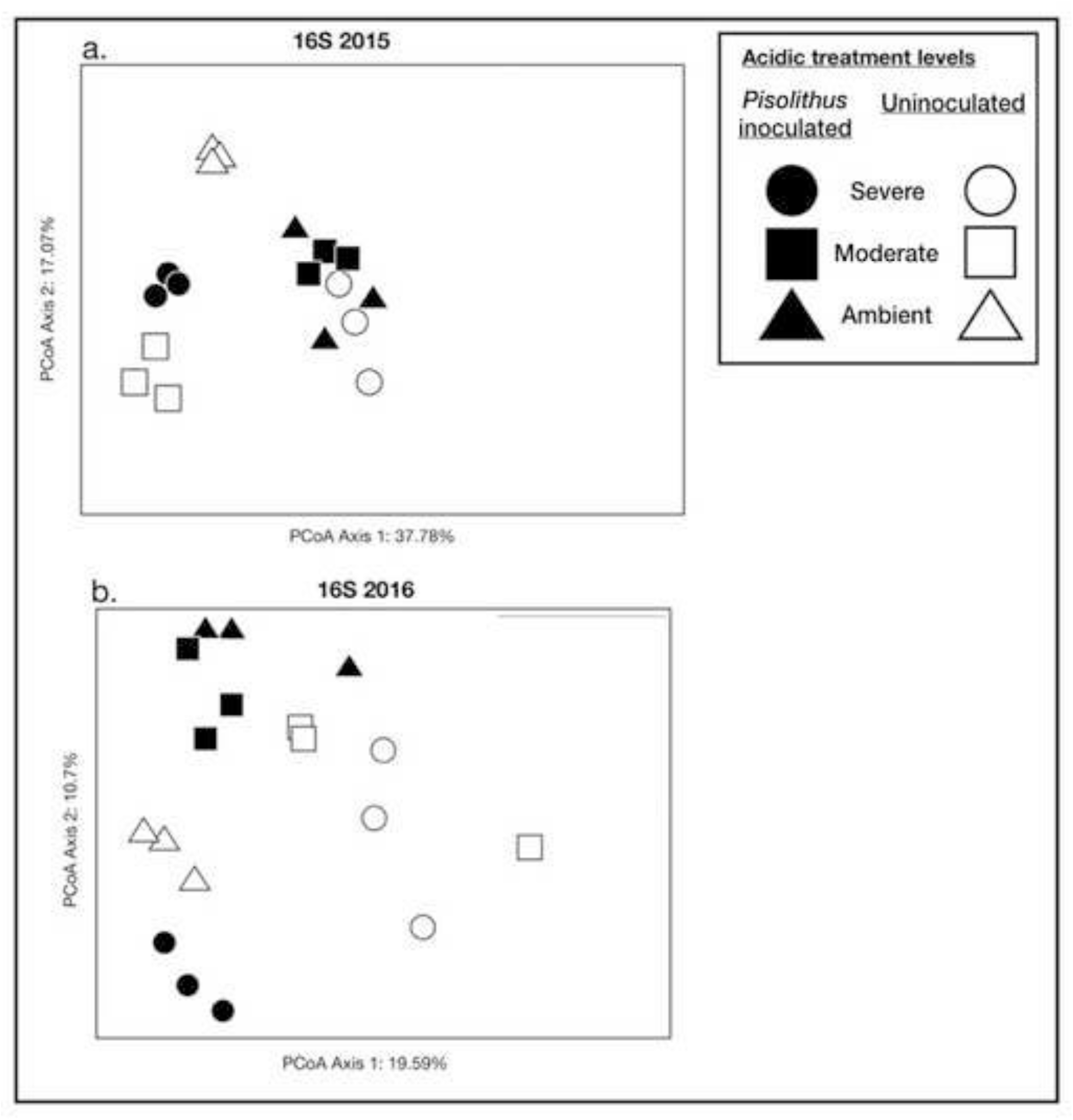
Principal coordinate analyses of bacterial communities. Principal coordinate analysis (PCoA) plot of 16S sequencing results from both 2015 (a) and 2016 (b), showing bacterial and archaeal community responses to acid and inoculation treatments. Solid shapes represent inoculation with *Pisolithus tinctorius*; open shapes indicate uninoculated treatments.

When we evaluated the effects of the ratio of divalent cations (Ca + Mg) to the total free Al concentrations on fungal community composition, we found that this ratio was related to fungal community composition in both 2015 (p=0.049) and 2016 (p = 0.001). For bacterial communities, we did not detect any significant effect of the ratio of divalent cations to the total free Al concentrations in 2015 (p=0.103) or for 2016 (p= 0.1449). However, in 2015 we detected a significant interaction between the pH of the treatment solutions and this ratio on bacterial communities (p= 0.001).

The PCoA Axis 1 in 2015 represented 37.78% of the variation in bacterial communities, while PCoA Axis 1 in 2016 represented 19.59%; Axis 2 in 2015 represented 17.07 %, and Axis 2 in 2016 represented 10.7% of the variation in bacterial communities (Fig 5). The most abundant bacterial order was Xanthomonadales, followed by Sphingobacteriales. In plots treated with severely-acid solutions, we found a greater relative abundance of Rhodocyclales and Clostridiales within the bacterial communities, as compared to the abundance of those particular orders found in plots treated with more moderate and less acidic treatment solutions.

Fungal functional groups varied in our treatment plots. There were greater numbers of fungal pathogens in plots inoculated with *Pisolithus,* then were found in uninoculated plots (p=0.015). Additionally, we detected greater saprotrophic fungal richness in plots treated with either moderately-acidic solutions (p=0.016) or severely-acidic solutions (p=0.030), than in ambient treatments plots. We found that acidic-solution treatment significantly affected the diversity of mycorrhizal fungal taxa (p<0.001), as did *Pisolithus* inoculation (p=0.047). We detected significant interactions between acidic-solution treatment and inoculation (p<0.001). We found that the ratio of divalent cations (Ca + Mg) to total free Al concentrations affected mycorrhizal fungal community composition (p=0.037), along with significant interactions between pH and this ratio on mycorrhizal fungal community composition (p=0.006), and a significant interaction among this ratio and *Pisolithus* inoculation on mycorrhizal communities (p=0.049).

The ectomycorrhizal taxon *Cenococcum* was found to dominate severely-acidic treatments inoculated with *Pisolithus*, and was barely detectable in the moderately-acidic treated or ambient treatments. In fact, *Cenococcum* was ∼ 200x more abundant in severely acidic inoculated treatment plants, than in all other treatments combined. Some EM fungal taxa, such as *Tricholoma* were only found in severely-acidic treated plots. While other EM taxa, such as *Russula, Sebacina,* and *Scleroderma* were only found in plots treated with moderately-acidic solutions. Another EM taxon, *Suillus,* was found only in ambient plots, and was not detected in either moderately or severely-acidic treatments. Fungal taxa *Amphinema* and *Archaeorhizomycetes* dominated ambient treatment plots, while *Trichophaea* and *Oidiodendron* were more prevalent in plots treated with severely-acidic solutions than in other treatment groups.

## Discussion and Conclusions

We examined the effects of long-term acidic treatments and mycorrhizal inoculation on above- and belowground properties in pine plantations in southern China. Overall, we found that severely-acidic treatments not only changed nutrient availability and organic matter content in soils, but also altered bacterial and fungal communities. Plant growth was inhibited and microbial community composition shifted, suggesting that long-term exposure to acidic treatments may affect both above- and belowground components of ecosystems. However, soils from plots inoculated with *Pisolithus* and exposed to severely-acidic treatments had very different fungal communities than those exposed to acid with autoclaved *Pisolithus* added. In the context of restoration, the results from these crossed treatments show that *Pisolithus* inoculation may alter fungal communities. In particular, inoculation may have elicited shifts towards fungal communities found in less disturbed soils or sites less impacted by acidic deposition, which may contain fungi that contribute towards stabilizing soil organic matter. Ostensibly, *Pisolithus* tolerates environmental stress and may either alleviate stressful conditions or prime soil microbial processes (Stroo and Alexander 1984, Silva et al. 2013). Although inoculation may influence the responses of microbes and some abiotic components of these terrestrial ecosystems to acid rain, benefits of *Pisolithus* inoculation were not evident for plants in severely-acidic treatments. Our findings highlight how biogeochemical cycling by plants and microbes may be constrained by long-term acidic deposition, but that inoculation with plant-associated mycorrhizal fungi may facilitate changes in the community structure of this forested ecosystem.

As terrestrial ecosystems are chronically exposed to acid rain, plants may become malnourished or their productivity may be diminished (Abdulaha-Al Baquy *et al.* 2017, Likens *et al.* 1996). Indeed, *Pinus massoniana* had lower needle biomass and plants were shorter in severely-acid treated plots, as compared to untreated ambient plots. As acidification inhibited plant growth and damaged aboveground tissues, it may have also hindered gas exchange and limited plant C sequestration (Kemmitt *et al.* 2006, Janssens *et al.* 2010, Oulehle *et al.* 2011). Although inoculation increased plant performance in moderately treated plots more than in ambient plots, inoculation did not significantly promote plant recovery in severely treated plots.

While the effects on fungal-plant mutualisms may be context-dependent, mycorrhizal associations may be expensive for plants to maintain in stressful environments (Jonsson *et al.* 2001, Hoeksema et al. 2010). EM fungi often augment plant primary production and plants reallocate a portion of this atmospheric C to their EM partner for use in building hyphae (Treseder and Holden 2013). However, environmental stress, such as strong acid, may hinder this process and halt both plant and fungal growth.

*Pisolithus* is a generalist melanized ectomyorrhizal fungus, which has been found to survive in highly acidic environments (Hendrix et al. 1985, Silva et al. 2013). As demonstrated by Fernandez et al. (2013) using minirhizotron imaging, fungal hyphae from another melanized EM fungal taxon, *Cenococcum geophilum*, may persist in the soil longer than non-melanized ectomycorrhizal fungi, as their melanin-rich hyphae is resistant to decomposition. Given that *Cenococcum* dominated our severely-acidic *Pisolithus* inoculated plots, and was barely detectable in any other treatments, the presence and persistence of this taxon may have contributed to our observed patterns of SOM accumulation, if these mycorrhizal residues contributed to soil C stocks (Clemmensen et al. 2013). Therefore, we posit that the prevalence of *Cenococcum* may have contributed to the marked increases in SOM found in our *Pisolithus* inoculated, severely-acidic treated plots.

Overall, we found lower SOM, available phosphorus, and nitrate in severely-acidic treated plots, possibly because the acid rain treatments modified the existing biogeochemical cycles. Acidic compound treatments may have stimulated microbes to use older and more recalcitrant carbon from soil organic carbon pools, as was found by Waldrop & Firestone (2004), which could contribute to these relatively low observations of SOM in severely-acidic, but uninoculated, treated plots. While the average acid status of the severely-acidic treated plots did not decrease at the end of the study period, the acidic water solution acidic inputs may have quickly and repeatedly dissolved nutrients from the soil, as was observed by Andersson *et al.* (2015). Because of its essential role in biomolecules and as a micronutrient, phosphorus is ubiquitous in soil biomass and critical to all living organisms. Fungal diversity has been correlated with higher plant-available phosphorus concentrations (Erlandson *et al.* 2016). However, since pH is a strong predictor of changes to fungal community composition, acidic stress could alter the fates of P during microbially-mediated organic matter turnover. Chemical weathering releases orthophosphate into soil solutions, yet, its bioavailability is optimized at neutral pH (pH ∼ 6.5) when Al and Ca phosphates are both minimized (Sylvia et al. 2005). Carrino-Kyker et al. (2016) found that elevating pH also increased soil P availability, while Barrow (1984) demonstrated P losses and reduced bioavailability at acidic pH levels.

The impact of acidic conditions on soil microbiota may have caused the variations we observed in nitrate availability. Although nitrate is mobile and readily assimilated by plants and microbes, acidic conditions may constrain the growth and activity of nitrifying bacteria (Sylvia et al. 2005). In our study, as the pH treatments approached neutral, we observed greater nitrate, available P, and SOM, than in severely-acidic treatment plots with autoclaved *Pisolithus* added. In contrast, we found no response of ammonia to acidification. Given the important role of mycorrhizal fungi in nitrogen cycling and supporting plant N acquisition, factors that constrain N cycling bacteria and fungi could influence the productivity of terrestrial ecosystems. Moreover, mycorrhizal fungi are sensitive to N availability; in an N deposition gradient in Alaska, Lilleskov et al. (2002) suggest that long-term N deposition dramatically changes ectomycorrhizal community structure and lead to reductions in ectomycorrhizal species richness. When *Pisolithus* inoculation treatments were applied to these plots treated with severely-acidic solutions, we observed an increase in available P, nitrate, and SOM, which suggests that *Pisolithus* may be assisting in biogeochemical cycling. Thus, adding a mycorrhizal fungus like *Pisolithus* may buffer soil conditions to promote nutrient retention, foster conditions conducive to co-existing taxa, or assist in the recovery of soil physicochemical properties in acidic conditions.

### Soil microbial communities

We found that the direct manipulations of simulated rain acidity extended beyond the plant community to affect soil microbes. *Pisolithus* inoculation significantly altered fungal and bacterial community composition. Further, the interactive effect of acidic treatment and inoculation status altered belowground fungal communities in a way that differed from the separate impact of each treatment.

Clemmenson (2015) showed that mycorrhizal functional types may strongly, and differentially, influence N immobilization and C sequestration in forested ecosystems. Therefore, the functional response of fungal communities to mycorrhizal inoculation may feedback to alter nutrient cycling or microbial interactions in ecosystems exposed to chronic acidic deposition. These findings highlight the potential of *Pisolithus* inoculum to buffer soil conditions for other fungi in ecosystems exposed to long-term acidic deposits.

Our study provides evidence that both acidic treatments and *Pisolithus* inoculation affect aboveground and belowground components of ecosystems, as well as soil physicochemical properties, yet it has some limitations. For instance, our study was conducted within pine plantations in southern China, and thus our interpretations are limited to *Pinus massoniana* forested ecosystems and plantations in this region. Additionally, although we applied acidic treatments to soils in our treatment plots weekly for 20 months, at the end of our study soil pH did not vary predictably across treatments. In fact, soil pH was as low in the acidic *Pisolithus* inoculated plots as it was in ambient plots with autoclaved *Pisolithus* added. Therefore, protracted additions of different levels of acidic solutions may have influenced the plant and microbial cycling of nutrients and water differently, which may have altered soil pH differently in our inoculated treatment plots, potentially because of stress-response traits associated with *Pisolithus*. Indeed, other studies have shown this fungus to be resilient and facilitate the recovery of plants grown in acidic coal spoils (Marx and Artman 1979).

It is worth noting that there was some inter-annual and plot-to-plot variation in soil pH, which in 2016, was significantly correlated with distance from the river. This may in turn have contributed to some of the effects and patterns observed in soil physicochemical properties or microbial communities throughout the study. However, we included distance from river as a factor in our statistical analyses, and did not detect effects of distance from river on initial soil pH or in 2015, and we have statistically proven differences driven by acidic-solution treatment and inoculation with *Pisolithus.*

Additionally, we attempted to prevent drought stress by applying additional water throughout the study. However, as the water used in between treatments had a normal pH for groundwater in the region (6.2), this may have contributed to the fact that the soil throughout the study did not show consistent acidifying effects of treatment, despite clear plant and microbial responses to the severely acidic treatment. Yet, weekly application of acidic solution treatments was applied for almost two years, which we found to affect both biotic communities and soil physicochemical properties. Overall, results from our acidic-solution treatments and inoculation manipulation suggest that *Pisolithus* inoculation may play a role in the remediation of forested ecosystems though facilitating changes to the mycorrhizal and microbial community, which could increase inputs into soil organic matter pools in chronically exposed severely-acidic ecosystems.

Our results provide evidence that soil abiotic and biotic components may be negatively affected by human-practices that increase acidic deposition, acid rain in particular. We found less SOM in severely-acidic treated plots; SOM, as illustrated by Magdoff and Weil (2004), may influence soil aggregate stability or improve plant growth. *Pisolithus* inoculation promoted both soil abiotic and fungal community recovery in severely-acidic treatment plots. Fungal community assembly in *Pisolithus* inoculated plots may include taxa which more efficiently use soil resources or are more effective at nutrient transfer to their plant hosts. However, severely-acidic inputs may damage aboveground vegetation and inhibit productivity, limiting C allocation to their associated EM partners. The resultant unequal bi-directional exchange of resources could feedback to promote fungal hyphal sloughing and the accumulation of C-rich fungal residues, which may contribute to our observed increases in soil organic matter in the *Pisolithus* inoculated severely-acidic treatment plots. As acidification directly reduces plant performance, it may also indirectly inhibit EM fungal proliferation.

Altogether, our findings suggest that chronic exposure to acidic inputs may change plant and soil properties and fungal and bacterial communities in exposed forested ecosystems. Moreover, inoculation with *Pisolithus* may assist in the recovery of some components of belowground ecosystems. These data have implications for management of acid rain-exposed forested ecosystems. For instance, restoration methods that aim to restore soil microbial communities, such as *Pisolithus* inoculation, may be especially effective at improving the soil nutrients and soil organic content at such acidified sites. This study underscores the importance of both reducing acidification in rain systems and maintaining soil microbial communities when restoring chronically exposed ecosystems.

## Acknowledgements

This research was supported by the Public Welfare Project of the National Scientific Research Institution (CAFRIFEEP201402) in China. We thank D. Jackson, G. Logan, B. Pickett, S. Saroa, M. Phillips, and A. Swanson for intellectual feedback and insightful comments on previous drafts. Moreover, we thank the Center for Conservation Biology and M.F. Allen and E.B. Allen for hosting ZC and MM. We are grateful for mycorrhizal fungal inoculum and inoculation support from the Study Center of Tree Mycorrhizae, from the Chinese Academy of Forestry. ZC was supported by the Public Welfare Project of the National Scientific Research Institution (CAFYBB2017SY026). EA, MM and KA were supported by the United States National Science Foundation (NSF ICER-1541047), EA was further supported by the National Science Foundation (NSF BIO-EF-1550920), and MM was supported by funding from the Mycological Society of America’s Translational Mycology Postdoctoral Award.

## Supplemental information

**Supplemental SI-Table 1.** Spatial arrangement of soil pH through time and by proximity to river. Spatial arrangement ascending in numbers with increasing distance from the Jingouba River.

| **Spatial block**<br>Proximity to River | **pH soil**<br>2015<br>Pre-treatment | **pH soil**<br>2015 | **pH soil**<br>2016 |
| --- | --- | --- | --- |
| **one** | 5.52 ± 0.37 | 5.69 ± 0.18 | 5.96 ± 0.34 |
| **two** | 5.65 ± 0.26 | 5.65 ± 0.18 | 6.31 ± 0.28 |
| **three** | 5.99 ± 0.48 | 5.72 ± 0.17 | 6.58 ± 0.20 |
| **four** | 5.85 ± 0.23 | 5.99 ± 0.13 | 6.71 ± 0.08 |

**Supplemental SI-Table 2.** Soil nutrients in our plots treated with acidic treatments and inoculated with *Pisolithus* in both 2015 and 2016.

| pH<br>solution<br>+/- *P.t.*<br>(*Pisolithus*<br>inoculation) | Available P<br>(mg kg<sup>-1</sup>) |  | Available<br>K<br>(mg kg<sup>-1</sup>) |  | NH<sub>4</sub>-N<br>(mg kg<sup>-1</sup>) |  | NO<sub>3</sub>-N<br>(mg kg<sup>-1</sup>) |  | Soil organic matter<br>(mg kg<sup>-1</sup>) |  |
| --- | --- | --- | --- | --- | --- | --- | --- | --- | --- | --- |
|  | 2015 | 2016 | 2015 | 2016 | 2015 | 2016 | 2015 | 2016 | 2015 | 2016 |
| Severe<br>3.5<br>+ *P.t.* | 9.0 ±<br>0.31 | 14.09 ±<br>0.76 | 193.33 ±<br>6.36 | 211.0 ±<br>7.09 | 7.92 ±<br>0.58 | 9.22 ±<br>2.01 | 239.78 ±<br>5.2 | 344.47 ±<br>39.87 | 17.75 ±<br>0.20 | 19.27 ±<br>0.29 |
| Severe<br>3.5 - | 6.0 ± 0.2 | 6.02 ±<br>0.18 | 160.33 ±<br>13.57 | 127.67 ±<br>2.91 | 1.97 ±<br>0.15 | 1.50 ±<br>1.50 | 118.54 ±<br>0.88 | 88.54 ±<br>6.31 | 11.82 ±<br>0.17 | 5.44 ±<br>0.25 |
| Moderate<br>4.5<br>+ *P.t.* | 9.83 ±<br>1.54 | 11.73 ±<br>0.76 | 163.67 ±<br>5.23 | 175.00 ±<br>4.04 | 3.73 ±<br>0.11 | 8.41 ±<br>2.40 | 153.78 ±<br>5.53 | 367.89 ±<br>23.16 | 13.22 ±<br>0.59 | 13.0 ±<br>0.01 |
| Moderate<br>4.5 - | 11.83 ±<br>1.17 | 15.57 ±<br>0.42 | 205.33 ±<br>12.17 | 259.33 ±<br>1.67 | 3.96 ±<br>0.39 | 7.38 ±<br>1.82 | 275.32 ±<br>4.08 | 708.19 ±<br>85.50 | 19.94 ±<br>0.65 | 17.60 ±<br>0.96 |
| Ambient<br>+ *P.t.* | 7.03 ±<br>0.28 | 9.51 ±<br>0.12 | 169.33 ±<br>0.67 | 200.0 ±<br>4.62 | 1.83 ±<br>0.22 | 4.06 ±<br>1.24 | 56.26 ±<br>3.35 | 227.35 ±<br>44.40 | 13.06 ±<br>0.27 | 13.79 ±<br>0.19 |
| Ambient<br>- | 14.45 ±<br>0.47 | 13.28 ±<br>0.18 | 207.67 ±<br>2.63 | 244.67 ±<br>1.33 | 3.58 ±<br>0.32 | 12.80 ±<br>1.35 | 180.32 ±<br>26.02 | 879.68 ±<br>34.12 | 20.05 ±<br>0.20 | 17.81 ±<br>0.83 |

**Supplementary Table.** Soil properties from 2015 in our plots treated with acidic treatments and inoculated with *Pisolithus*. Elemental analyses included sodium (Na), calcium (Ca), magnesium (Mg), and aluminum (Al), as well as cation exchange capacity (CEC).

| pH solution<br>+/- *P.t.*<br>(*Pisolithus*<br>inoculation) | Na<br>(cmol kg<sup>-1</sup>) | Ca<br>(cmol kg<sup>-1</sup>) | Mg<br>(cmol kg<sup>-1</sup>) | Al<br>(mg kg<sup>-1</sup>) | CEC<br>(cmol kg<sup>-1</sup>) |
| --- | --- | --- | --- | --- | --- |
| Severe<br>3.5 + *P.t.* | 1.0 ± 0.06 | 6.45 ± 0.14 | 0.42 ± 0.02 | 7.02 ± 2.19 | 7.08 ± 0.98 |
| Severe<br>3.5 - | 0.75 ± 0.04 | 5.57 ± 0.16 | 0.95 ± 0.05 | 2.12 ± 0.27 | 7.69 ± 0.08 |
| Moderate<br>4.5 + *P.t.* | 0.67 ± 0.02 | 4.42 ± 0.07 | 1.25 ± 0.36 | 3.23 ± 0.29 | 8.54 ± 0.43 |
| Moderate<br>4.5 - | 0.85 ± 0.11 | 8.22 ± 0.97 | 0.44 ± 0.01 | 1.66 ± 0.09 | 8.49 ± 0.41 |
| Ambient<br>+ *P.t.* | 0.73 ± 0.03 | 4.95 ± 0.09 | 0.57 ± 0.01 | 2.52 ± 0.99 | 8.3 ± 1.35 |
| Ambient - | 0.61 ± 0.01 | 5.12 ± 0.70 | 0.39 ± 0.01 | 6.34 ± 1.3 | 7.76 ± 0.77 |

**Supplemental SI-Table 4.** Bacterial community diversity metrics from 2016. Metrics include: Abundance-based Coverage Estimate (ACE), Chao1, Fisher, Observed OTUs, Shannon, and Simpsons.

| pH<br>solution<br>+/- *P.t.*<br>(*Pisolithus*<br>inoculation) | ACE | Chao1 | Fisher | Observed<br>OTUs | Shannon | Simpson |
| --- | --- | --- | --- | --- | --- | --- |
| 3.5 + *P.t.* | 2642.04 ±<br>24.75 | 2639.84 ±<br>50.80 | 601.62±<br>12.03 | 2178.67±<br>31.84 | 6.57±<br>0.019 | 0.995±<br>0.0003 |
| 3.5 – | 2587.61 ±<br>24.06 | 2625.95 ±<br>51.97 | 594.80±<br>26.17 | 2159.67±<br>70.11 | 6.53±<br>0.069 | 0.997±<br>0.0004 |
| 4.5 + *P.t.* | 2868.45 ±<br>25.69 | 2883.93 ±<br>54.57 | 672.13±<br>11.16 | 2361.67±<br>28.48 | 6.73±<br>0.028 | 0.997±<br>0.0001 |
| 4.5 – | 2793.60 ±<br>25.45 | 2796.59±<br>52.2 | 652.28±<br>37.44 | 2309±<br>97.22 | 6.74±<br>0.113 | 0.997±<br>0.0006 |
| 5.6 + *P.t.*<br>(Ambient) | 2875.35 ±<br>25.04 | 2887.16 ±<br>57.23 | 662.43±<br>23.08 | 2336.33±<br>63.38 | 6.70±<br>0.046 | 0.997±<br>0.0001 |
| 5.6 –<br>(Ambient) | 2696.85 ±<br>25.31 | 2694.06 ±<br>55.28 | 600.43±<br>5.96 | 2175.67±<br>15.76 | 6.48±<br>0.025 | 0.995±<br>0.0004 |

**Supplemental SI-Table 5.** Fungal community diversity metrics from 2016. Metrics include: Abundance-based Coverage Estimate (ACE), Chao1, Fisher, Observed OTUs, Shannon, and Simpsons.

| pH<br>solution<br>+/- *P.t.*<br>(*Pisolithus*<br>inoculation) | ACE | Chao1 | Fisher | Observed<br>OTUs | Shannon | Simpson |
| --- | --- | --- | --- | --- | --- | --- |
| 3.5 + *P.t.* | 480.39 ±<br>20.43 | 477.28 ±<br>16.31 | 63.31±<br>3.42 | 389.0±<br>17.06 | 3.03±<br>0.079 | 0.84±<br>0.006 |
| 3.5 – | 439.98 ±<br>8.91 | 451.29 ±<br>7.95 | 56.54±<br>0.88 | 353.67±<br>4.48 | 2.97± 0.2 | 0.89±<br>0.013 |
| 4.5 + *P.t.* | 538.29 ±<br>29.61 | 541.03 ±<br>37.33 | 75.36±<br>2.06 | 450.0±<br>10.26 | 3.73± 0.04 | 0.94±<br>0.005 |
| 4.5 – | 496.21 ±<br>18.22 | 475.001±<br>17.34 | 67.94±<br>2.70 | 412.67±<br>13.67 | 3.35± 0.02 | 0.90±<br>0.011 |
| 5.6 + *P.t.*<br>(Ambient) | 540.38 ±<br>22.99 | 523.13 ±<br>25.7 | 75.58±<br>2.81 | 451.0±<br>14.47 | 3.58± 0.09 | 0.93± 0.01 |
| 5.6 –<br>(Ambient) | 429.08 ±<br>17.88 | 428.87 ±<br>14.51 | 65.11±<br>3.35 | 398.0±<br>17.57 | 3.63± 0.12 | 0.92±<br>0.011 |

**Supplemental Figure 1.**
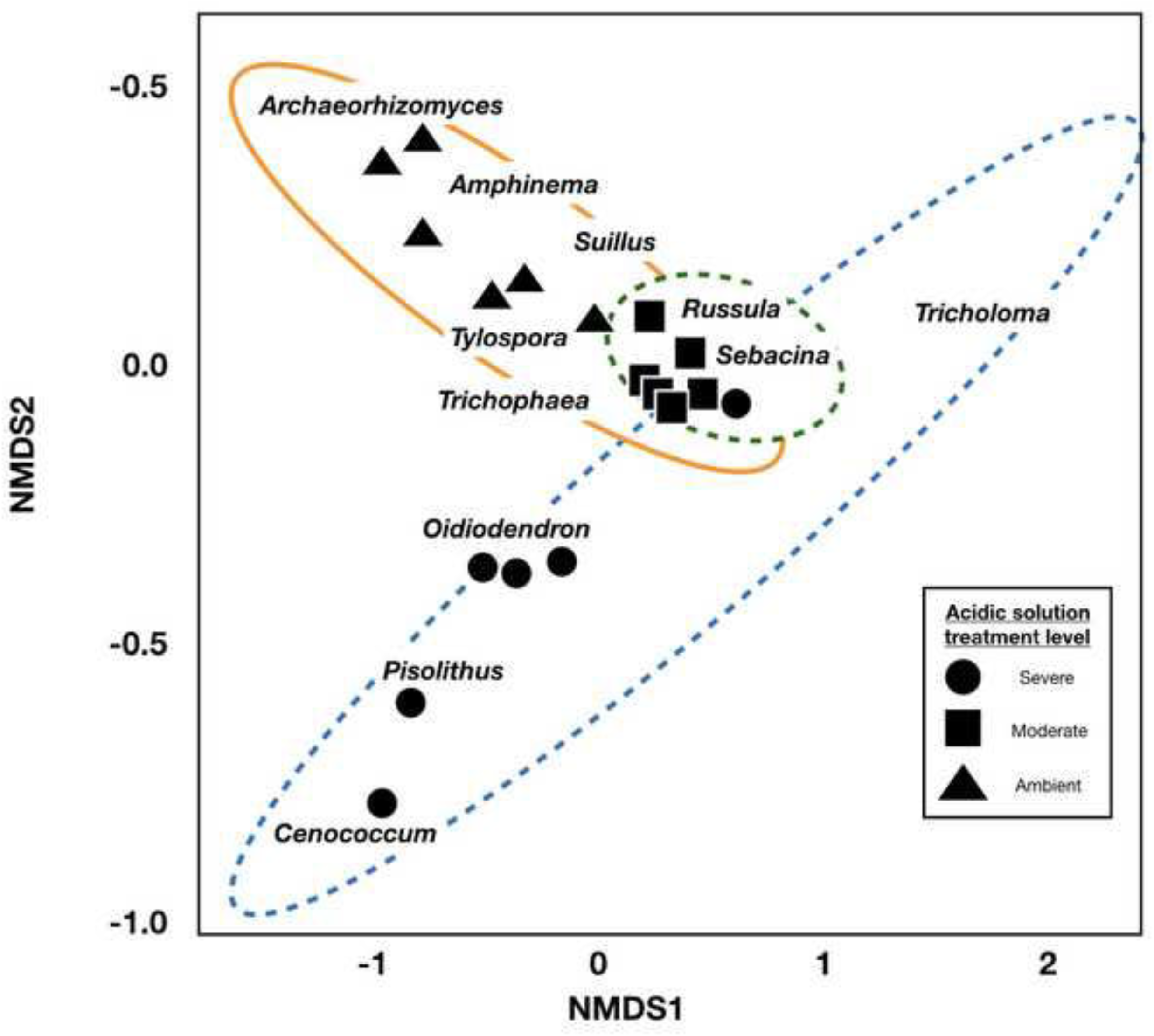
Non-metric multi-dimensional scaling plot illustrating differences in key fungal taxa by acidic-solution treatment. Shapes represent a portion of the fungal community from either severely acidic treatment plots (circles) shown surrounded by blue dashed ellipse, moderately acidic treatments (squares) shown by the green dashed ellipse, or ambient treatments (triangles) represented by the orange solid ellipse. Ectomycorrhizal fungal taxa (e.g., *Suillus*) and putative root endophytes (e.g., *Archaeorhizomycetes*) associated with particular acidic treatment categories are shown in text in close proximity to that representative ellipse.

**Supplemental Figure 2.**
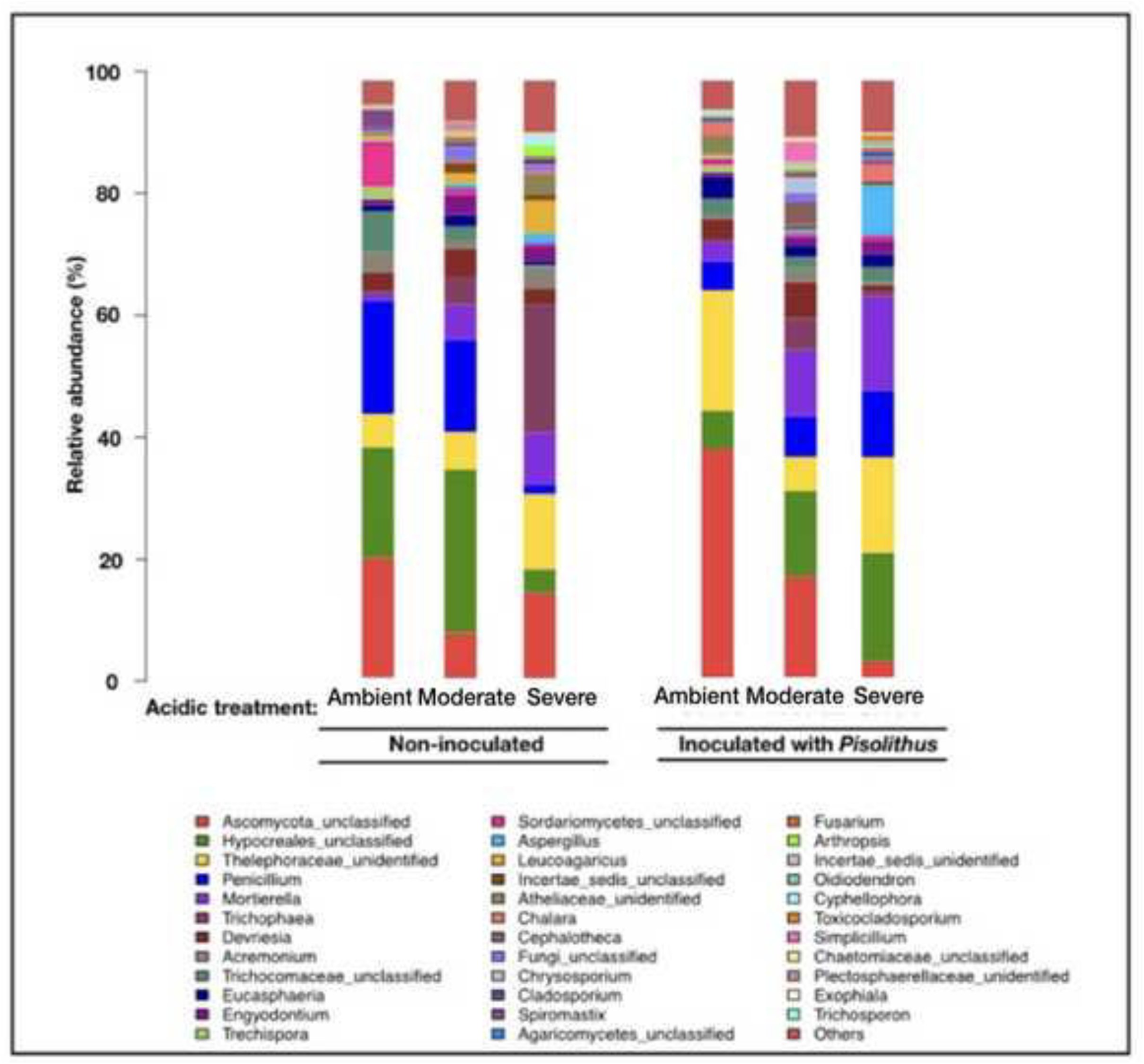
Fungal taxa plotted in bar charts, shown with the relative abundance of fungal taxa by acidic-solution treatment and by *Pisolithus* inoculation; treatments with autoclaved *Pisolithus* added marked as ‘Non-inoculated.’

**Supplemental Figure 3.**
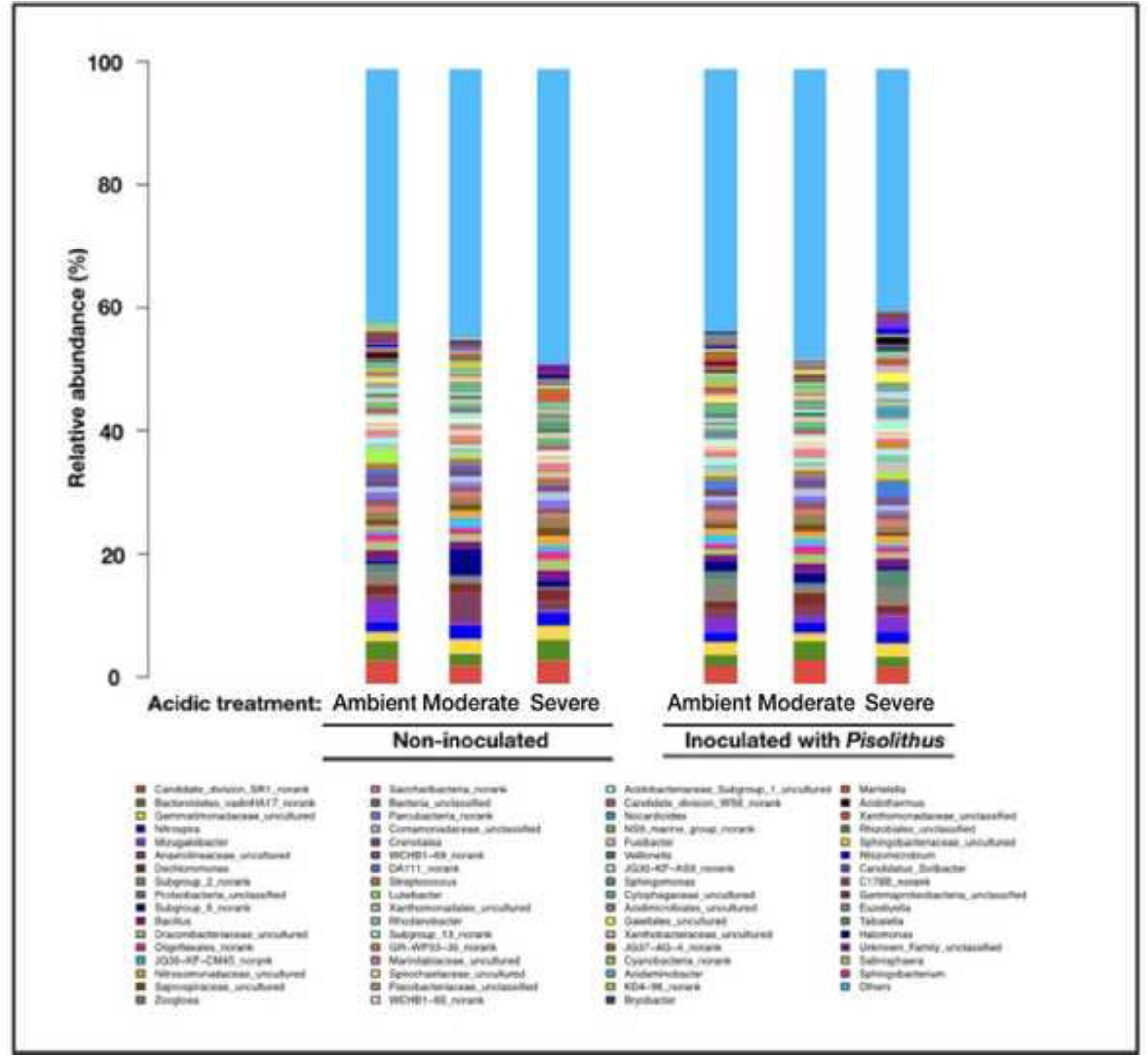
Bacterial taxa plotted in bar charts, shown with the relative abundance of fungal taxa by acidic-solution treatment and by *Pisolithus* inoculation; treatments with autoclaved *Pisolithus* added marked as ‘Non-inoculated.’

**Supplemental Methods:** Molecular methods used on our soil microbial analyses.

## Supplemental molecular methods: SI

We extracted total DNA from each of the three 0.25 g soil aliquots using an E.Z.N.A.^®^ soil DNA Kit (Omega Bio-tek, Norcross, GA, U.S.) according to manufacturer’s protocols. We pooled these DNA extracts into a single representative DNA extract and subsequently standardized concentrations to 10 ng/µl before Polymerase chain reactions (PCR) amplification. We targeted hypervariable portions of both the V3–V4 region of the bacterial 16S rRNA gene and the fungal internal transcribed spacer (fungal ITS1) region using primers modified for Illumina Miseq sequencing (Illumina Incorporated, California).

For bacteria, the V3-V4 region of 16S was amplified using forward primer 338F (5’-ACTCCTACGGGAGGCAGCAG-3) and reverse primer 806R (5’-GGACTACHVGGGTWTCTAAT-3’), along with a unique eight-base barcode assigned to each sample. For fungi, the internal transcribed spacer region was amplified with forward primer ITS1 (5’ - CTTGGTCATTTAGAGGAAGTAA-3’) and reverse primer 2043R (5’-GCTGCGTTCTTCATCGATGC-3’). Fungal ITS1 region was targeted, along with unique molecular barcodes assigned to each reverse primer. For both 16S and ITS1, polymerase chain reactions were set up in triplicate using 0.8 µl of each primer (5 µM), 0.4 µL of FastPfu Polymerase, 10 ng of DNA template, 2 µL of 2.5 mM dNTPs, 4 µL of 5 × FastPfu Buffer. Reactions ran for 95 °C for 3 min, followed by 25 cycles at 95 °C for 30 s, 55 °C for 30 s, and 72 °C for 30 s and a final extension at 72 °C for 10 min.

Both 16S and ITS, PCR amplicons were visualized using gel electrophoresis, and subsequently extracted from 2% agarose gels. Amplicons were purified using a AxyPrep DNA Gel Extraction Kit (Axygen Biosciences, Union City, California) according to the manufacturer’s instructions, and then quantified in individual tubes using QuantiFluor™ -ST (Promega, Madison, Wisconsin). Purified amplicons were pooled in equimolar concentrations for downstream Illumina sequencing.

Our PCR products were sequenced on an Illumina MiSeq flowcell by Shanghai Majorbio (Shanghai, China) and paired-end sequences were generated (2 × 300) according to the standard protocols. Raw fastq files were demultiplexed, quality-filtered using QIIME (version 1.17); 300 basepair (bp) reads were truncated at quality scores <20 over a 50 bp range and truncated reads shorter than 50bp were discarded. Barcode matching allowed for any two nucleotide base mismatches, however, reads containing ambiguous characters were removed. Only sequences overlapping longer than 10 bp were assembled according to their overlap sequence. Operational Units (OTUs) were clustered with 97% similarity cutoff using UPARSE (version 7.1 http://drive5.com/uparse/) (Edgar 2013). Chimeric sequences were identified and removed using UCHIME. Taxonomic information associated with each 16S rRNA gene sequence was analyzed by RDP Classifier (http://rdp.cme.msu.edu/) against the Silva (SSU115) 16S rRNA database and taxonomy of each ITS1 gene sequence was analyzed by RDP Classifier (http://rdp.cme.msu.edu/) against the UNITE 7.0 database.

